# Peptide recognition and dephosphorylation by the vaccinia VH1 phosphatase

**DOI:** 10.1101/2020.05.26.100743

**Authors:** Bryan M. Zhao, Megan Hogan, Michael S Lee, Beverly K. Dyas, Robert G. Ulrich

## Abstract

The VH1 protein encoded by the highly conserved H1 locus of orthopoxviruses is a dual-specificity phosphatase (DUSPs) that hydrolyzes phosphate groups from phosphorylated tyrosine, serine, and threonine residues of viral and host cell proteins. Because the DUSP activities are required for virus replication, VH1 is a prime target for the development of therapeutic inhibitors. However, the presentation of a shallow catalytic site has thwarted all drug development efforts. As an alternative to direct targeting of catalytic pockets, we describe surface contacts between VH1 and substrates that are essential for full activity and provide a new pathway for developing inhibitors of protein-protein interactions. Critical amino acid residues were manipulated by site-directed mutagenesis of VH1, and perturbation of peptide substrate interactions based on these mutations were assessed by high-throughput assays that employed surface plasmon resonance and phosphatase activities. Two positively-charged residues (Lys-20 and Lys-22) and the hydrophobic side chain of Met-60 appear to orient the polarity of the pTyr peptide on the VH1 surface, while additional amino acid residues that flank the catalytic site contribute to substrate recognition and productive dephosphorylation. We propose that the enzyme-substrate contact residues described here may serve as molecular targets for the development of inhibitors that specifically block VH1 catalytic activity and thus poxvirus replication.

## INTRODUCTION

Poxviruses (*Poxviridae*) are large double-stranded DNA viruses that replicate exclusively in the cytoplasm of vertebrate and invertebrate host cells. Notable orthopoxviruses within the *Poxviridae* family include monkeypox viruses circulating in Central Africa that have 10% case-fatality rates (1), the variola major virus of smallpox that carries a 30% fatality rate within non-vaccinated populations (2), posing a threat as a bioterrorist weapon, and vaccinia virus strains used as vaccines to control monkeypox and smallpox infections. The orthopoxvirus genome encodes kinases B1 and F10 that phosphorylate six or more virus-encoded proteins (3–5), as well as a dual-specificity protein phosphatase (DUSP) VH1 that removes phosphate groups from phosphorylated Ser (pSer), Thr (pThr) and Tyr (pTyr) residues of protein substrates (6). The kinases and VH1 phosphatase are essential for viral replication (7). Significantly, phosphorylation of tyrosine residues contained within host proteins is also controlled by the reversible actions of cellular protein tyrosine kinases (PTKs) and protein tyrosine phosphatases (PTPs), including the DUSPs (8,9). These phospho-transfer enzymes critically affect cell proliferation, metabolism, adhesion, cell growth, and cell differentiation (10–13). VH1 was the first DUSP described (14), and is highly conserved within the orthopoxvirus genus (Figure 1). Active sites of VH1 phosphatases share with most DUSPs a highly conserved PTP-loop (HCXXXXXR) that harbors the catalytic thiol of a nucleophilic Cys residue and a general acid residue (Asp). Genetic repression of VH1 expression leads to the production of noninfectious virions (7), and studies suggest that VH1 overcomes host defense mechanisms during infection by dephosphorylating the Signal Transducer And Activator Of Transcription 1 (STAT1), thereby blocking the interferon-γ signaling pathway (15). Because VH1 does not dephosphorylate DNA bound STAT1, the viral phosphatase may act predominantly on the cytoplasmic pool of activated STAT1 (16).

**Figure 1.**
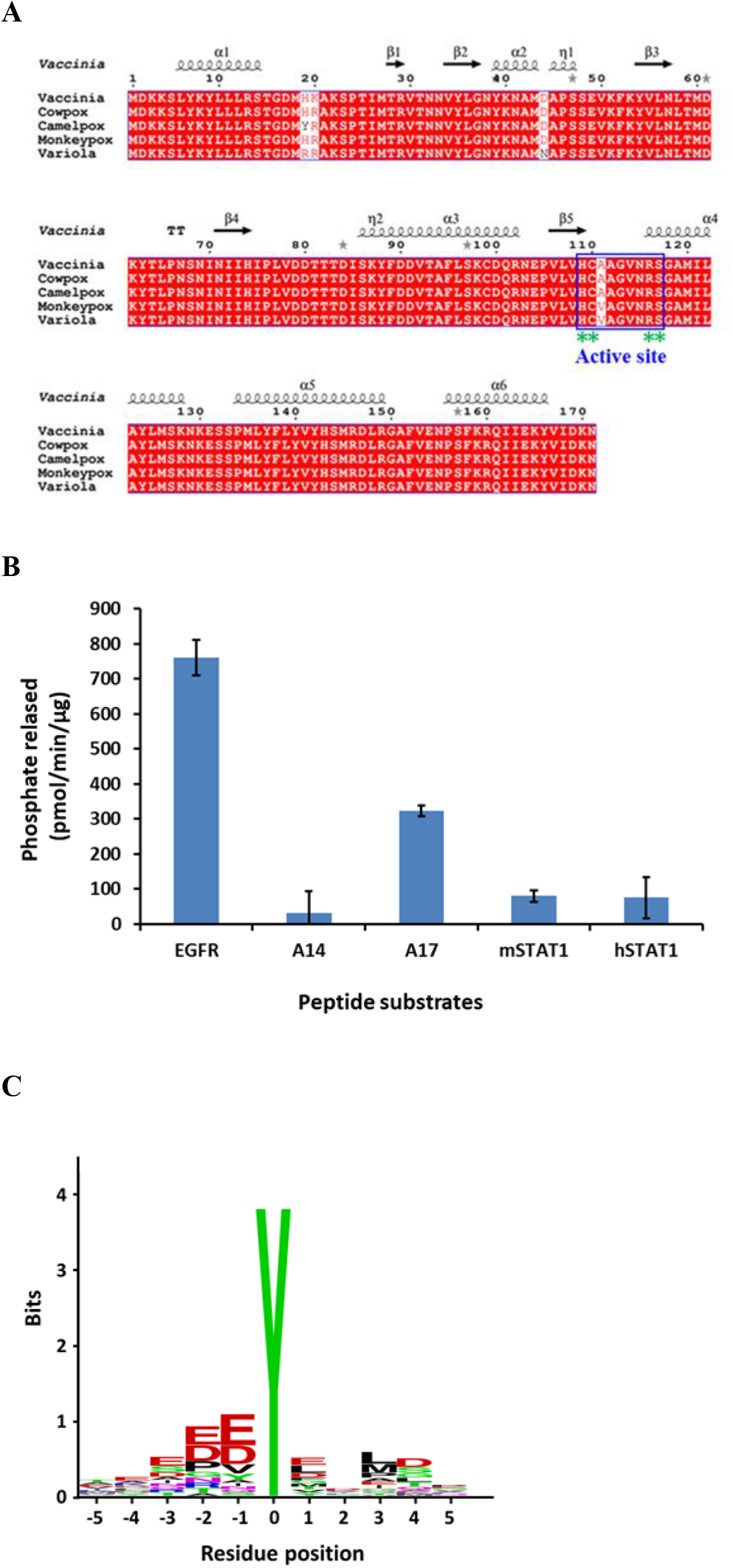
Sequence motifs for the VH1 catalytic site and phosphorylated peptide substrates. **A.** Multiple sequence alignment of H1 phosphatases from orthopoxviruses. The conserved residues are highlighted in red. The blue box indicates the catalytic site of VH1 proteins with conserved residues indicate by green asterisks. The number above the sequences corresponds to positions of VH1 residues. The secondary structure elements were generated based on the vaccinia VH1 structure (PDB: 3CM3), and strands are indicated by arrows and coils for helices. The figure for multiple alignments was generated using ESPript 3.0 (43). VH1 catalytic site sequences from diverse poxvirus are colored in yellow. Sequence variations (white background) occur at the amino acid position 19, 20, 44 and 111. **B.** VH1 dephosphorylation of select viral and host peptide substrate. Released phosphate was quantified by using a malachite green assay. **C.** Consensus peptide substrate sequence from library screening. Bits represent the relative frequency of amino acids based on the top 30 phosphorylated peptides.

While catalytic sites of many protein tyrosine phosphatases (PTPs) are challenging for drug discovery efforts, a strategy of targeting secondary binding pockets along with active site residues has resulted in significant progress in the development of highly potent and selective inhibitors (17–19). However, the crystal structure of VH1 reveals a much shallower catalytic cleft (6 Å) compared to other PTPs, for example, PTP1B (9 Å)(16,20), which is more difficult for the design of small-molecule inhibitors. The study described here examines the protein-protein interactions of VH1 as a possible alternative guide for inhibitor development. By screening a library of pTyr peptides from host cell proteins, we identified a pTyr-peptide of EGFR as a highly potent substrate and evaluated the optimal peptide length for VH1 catalytic activity (*K_m_*) in comparison with binding (*K_D_*). Amino acids that mapped to protein surfaces around the catalytic site of VH1 were sequentially mutated to alanine to examine the effects on catalytic activity, peptide binding, and VH1 thermal stability (*T*_m_). Peptide substrate binding and dephosphorylation were evaluated *en masse* by using unique protein microarray assays. Finally, based on the experimental results and known protein structures, a model of the peptide binding footprint on VH1 was developed to guide future development of inhibitors.

## RESULTS

### VH1 peptide phosphatase assay

The vaccinia VH1 phosphatase was expressed in *Escherichia coli* as a full-length, 187-amino acid protein (18-kDa) and a fusion of VH1 to maltose-binding protein (MBP)-poly His tag to increase protein expression levels (21). The recombinant proteins were purified by Immobilized Metal Affinity Chromatography (IMAC) as previously described (supplemental Figure 1) (22). The activities of both native VH1 and MBP-VH1 fusion proteins were confirmed by hydrolysis of the small molecular substrate, *para*-nitrophenylphosphate (pNPP). We next investigated enzymatic activity with phosphorylated peptides. Immune responses that are mediated by STAT1 are inhibited by VH1 (15), while VH1-dependent dephosphorylation of the poxvirus proteins A17 and A14 is necessary for maturation and assembly of infection-competent viruses (23). However, these intracellular protein substrates for VH1 were previously identified primarily by cell-culture assays (15,23), and little information is available for the *in vitro* activity of peptides based on the essential sequences for DUSP recognition. Therefore, we synthesized pTyr peptides from human STAT1(hSTAT1), mouse STAT1 (mSTAT1), and vaccinia A17, along with an A14 pSer peptide from A14 (Table1), and examined *in vitro* activity of VH1 by liberation of phosphate. Although all four peptides were dephosphorylated by VH1, the observed reaction rates were too slow (Figure 1B) to allow us to establish a robust assay. To identify peptide substrates that had greater activity with VH1 compared to the putative endogenous cellular and viral proteins, we screened a phosphorylated peptide substrate library that contained a selection of 360 pTyr-peptide sequences that were derived from phosphorylation sites of human cellular proteins. A consensus amino acid sequence of the most active peptides from the phosphorylated peptide library screening (Figure 1C) included dominant negatively charged residues around the pTyr residue. These results with vaccinia virus VH1 were consistent with previous data for VH1 from variola virus (24) that demonstrated the requirement for negatively charged amino acid residues near the pTyr. Comparing these two viral DUSPs, there are four amino acid sequence differences between vaccinia and variola virus VH1, as shown in Figure 1A, while the active site is completely conserved among the orthopoxviruses. Among the most active VH1 peptide substrates from the library screening, an EGFR pTyr peptide ^986-^VDADEpYLIPQQ^-998^ was selected for further evaluation. As expected, the variola VH1 dephosphorylated the EGFR peptide with activity levels that were equal to vaccinia VH1 (data not shown). Interestingly, the EGFR peptide identified by these results is also a potent PTP peptide substrate for YopH and PTP1B (25–27).

**Table1.** Phospho-Tyrosine peptides used to study VH1 activity.

| Peptide Name | Amino acid sequences | pTyr/Ser | Protein name | GenBank™ Accession# |
| --- | --- | --- | --- | --- |
| hSTAT1 | <sup>695</sup>-GPKGTG(**pY**)IKTELISVSEV<sup>-701</sup> | 701 | Human STAT1 | ADA59516 |
| mSTAT1 | <sup>693</sup>-LDDPKRTG(**pY**)IKETLISVS<sup>-702</sup> | 702 | Mouse STAT1 | AAA19454 |
| A14 | <sup>78</sup>-GVIHTNH(**pS**)DISMN<sup>-90</sup> | 85 | Vaccinia A14 | AND73968 |
| A17 | <sup>193</sup>-PTFNSLNTDD(**pY**)<sup>-203</sup> | 203 | Vaccine A17 | P68593 |
| EGFR 11mer | <sup>987</sup>-VDADE(**pY**)LIPQQ<sup>-997</sup> | 992 | EGFR | NP_005219 |
| EGFR 8-mer | <sup>987</sup>-VDADE(**pY**)LI<sup>-994</sup> | 992 | EGFR | NP_005219 |
| EGFR 3-mer | <sup>991</sup>-E(**pY**)L<sup>-993</sup> | 992 | EGFR | NP_005219 |

We varied the length of the EGFR peptide to determine if amino acid residues that flank the pTyr impacted interactions with VH1. As shown in Table 2, the EGFR 11-mer and 8-mer peptides have similar *K_m_* values, and the *K_m_* for a 3-mer peptide was reduced only by 50%. These results suggested that reducing the length of the EGFR peptide had little direct effect on enzymatic activity. We generated a substrate-trapping mutant of VH1 (mVH1) to determine the binding interactions of the EGFR peptides with VH1. By replacing the active site Cys-110 with Ala, the substrate-trapping mutant mVH1 was catalytically inactive to allow us to measure peptide binding by surface plasmon resonance (SPR). The *K*_D_ values for the 11-mer, 8-mer, and 3-mer EGFR pTyr peptides were determined to be 13μM, 335 μM, and 1880 μM, respectively (Table 2). We concluded that although the EGFR 11-mer and 8-mer peptides have similar *K_m_* values, the *K*_D_ values differed by 20-fold. This striking difference in affinity compared to catalytic activity among the different length EGFR peptides strongly suggested the importance of residues flanking pTyr in stabilizing peptide binding to VH1.

**Table 2.** Effect of phosphorylated peptide length on VH1 catalysis and substrate binding.

| | | $K_D$ (μM)* | $K_m$ (μM) ** |
| --- | --- | --- | --- |
| EGFR 11-mer | Bio-ahx-VDADE(pY)LIPQQ | 13 | 257 |
| EGFR 8-mer | Bio-ahx-DADE(pY)LI | 335 | 226 |
| EGFR 3-mer | Bio-ahx-E(pY)L | 1880 | 537 |

\*Determined by SPR with immobilized peptides and soluble VH1 C110S.
\*\*VH1 wild type was incubated with 1-300 μM peptide for 15 min, 37 °C. The amount of phosphate released was determined by comparison to a standard curve. $K_m$ values were expressed as the mean ± S.D. (triplicate experiments).

### VH1-peptide interaction surfaces determined by catalytic activity

To gain a more detailed understanding of the EGFR peptide interactions with VH1, we sequentially replaced 25 protein surface residues around the shallow catalytic pocket of VH1 and examined the effect of each alanine substitution on the catalytic activity of VH1 protein (VH1-MBP fusion) with the small molecule substrate 6,8-difluoro-4-methylumbelliferyl phosphate (DiFMUP) and the EGFR peptide (Figure 2A). We hypothesized that the alanine substitutions should have a significant impact on EGFR peptide dephosphorylation if a residue was involved in stabilizing peptide binding, and have minimal effect on the hydrolysis of the small molecule substrate DIFMUP (molecular weight: 292.13226). VH1 mutants that exhibited a significant reduction in catalytic activity in the DiFMUP assay, regardless of the effect on peptide dephosphorylation, were eliminated from further consideration. These included residues Asn-38, Tyr-39, Lys-40, Asp-44, Asp-61, Asp-79, A112, Val-114, Tyr-142, Asn-155, Glu-154, and Lys-159 (Figure 2B, C). Conceivably, reduced DiFMUP activity resulted from direct perturbation of the VH1 catalytic site or destabilizing effects on the overall protein structure of VH1. We also noted residues that could be substituted with only negligible decreases in catalytic activity towards the EGFR peptide or DiFMUP: His-19, Lys-20, Lys-22, Ser-23, Thr-25, Ile-26, Thr-81A, Thr-82 and A111V (Figure 2 B, C). Most importantly, we identified mutated residues of VH1 (Met-18, Ala-21, Thr-59, Met-60, Val-78 and Asp-80) that presented DiFMUP hydrolysis levels that were similar to wild-type VH1 but with reduced EGFR dephosphorylation, and concluded these residues were most likely to be linked to interactions between peptide and VH1 (Figure 2B, C). We further noted that mutations in amino acid residues Tyr-142, Glu-154, and Lys-159 greatly diminished phosphatase activity possibly due to disruption of the previously described dimer formation (28) that is important for optimal phosphatase activity.

**Figure 2.**
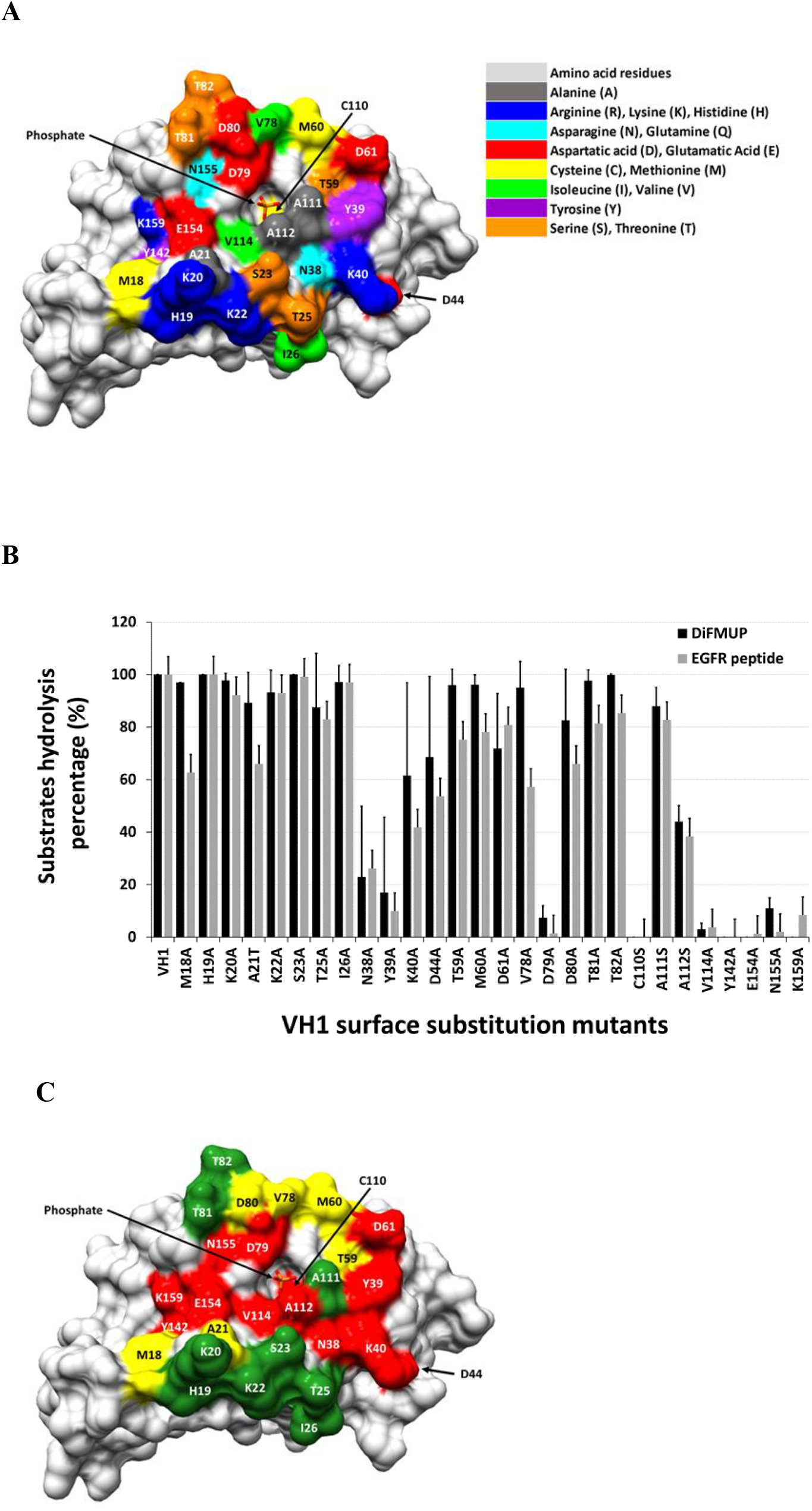
Molecular surfaces of VH1 linked to pTyr-peptide dephosphorylation. **A.** Molecular surface representation of residues surrounding the VH1 catalytic site that were examined in this study. The selected residues are colored according to chemical properties. **B.** Phosphatase activities of the VH1 mutants for small molecule or phosphorylated peptide substrates. Dephosphorylation results for each alanine substitution mutant of VH1 were normalized to the wild-type VH1. Error bars indicate ± SD for three independent experiments. **C.** Protein surface representation of VH1 site-specific mutagenesis results. Inhibition of DiFMUP hydrolysis, red-colored residues; inhibition of only phosphorylated peptide hydrolysis, yellow-colored residues; and no effect in either assay, green-colored residues.

### Structural stability of VH1 protein surface mutants

We sought to determine if any of the mutations introduced into VH1 could have destabilized protein structure or, conversely, if any of the possible protein-substrate contact residues contributed to protein stability. A thermal shift assay was used to determine changes in the melting temperature (Δ*T*_m_) for each of the VH1 mutant proteins under identical experimental conditions. As shown in Figure 3A, the calculated *T*_m_ value for the wild-type VH1 (MBP-VH1) was 54 °C, while the catalytically inactive C110S mutant was the most stable of all VH1 that we examined, with a *T*_m_ that was shifted by two degrees (56 °C). We calculated the standard deviation for all mutants of VH1 and used the 1.5 standard deviations (1.5 °C) above or below the mean as cut off value. Most of the VH1 site-specific mutants proteins had slightly reduced *T*_m_ values (not less than −1.5 °C) compared to the wild-type VH1 (Figure 3B), which represented 86.7% of all mutant VH1. In contrast to C110S, mutations of residues Tyr-39, Lys-40, Asp-61, and Lys-159 resulted in the least stable structures, with Δ*T*_m_ values shifted by −1.5 to −3 °C.

**Figure 3.**
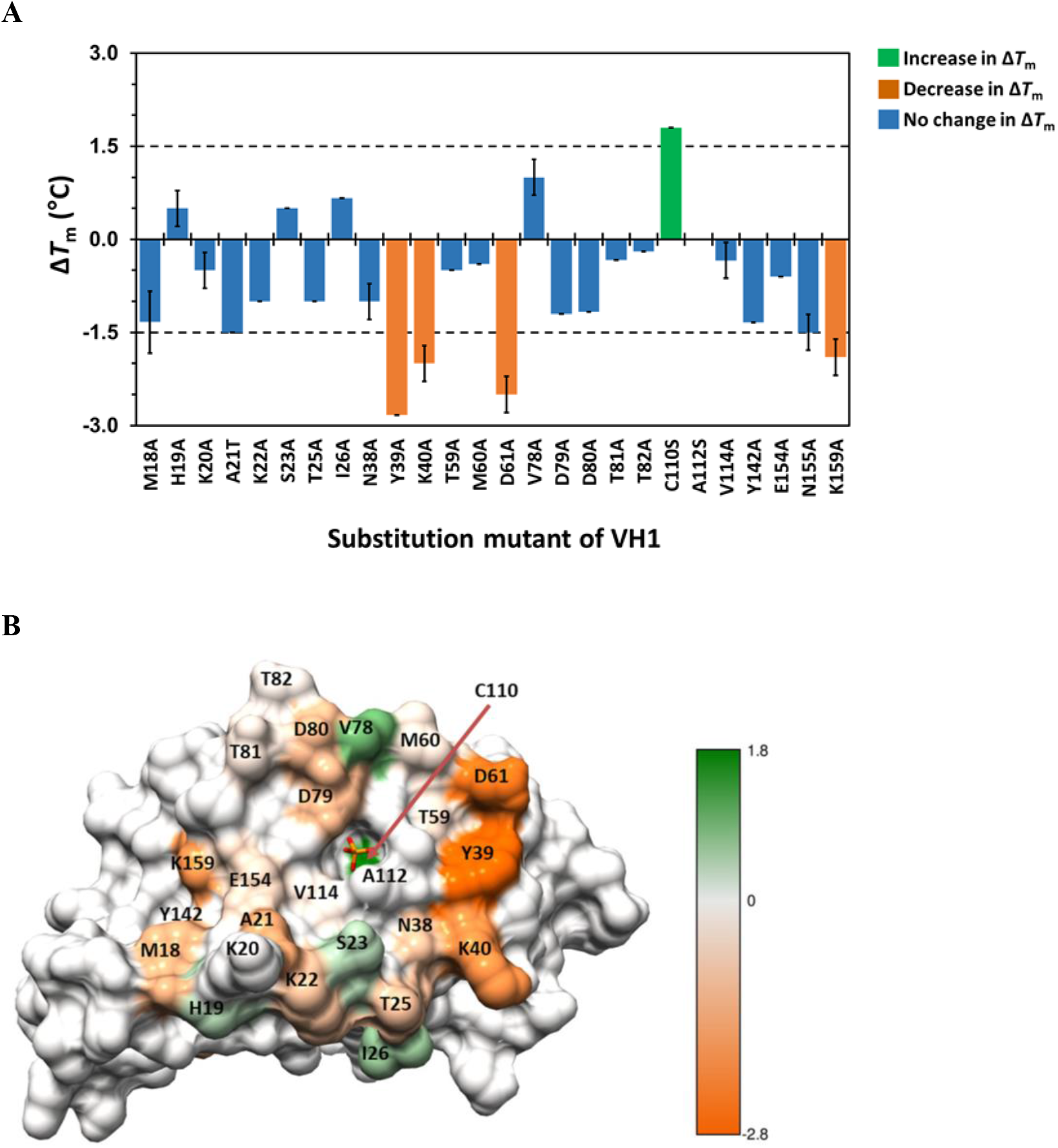
Thermostability of VH1 mutants. **A.** Δ*T*_m_ values for mutated VH1 proteins. The horizontal dotted lines indicate the cut-off values (±1.5 SD). **B.** Surface representations of VH1 phosphatase with residues colored by Δ*T*_m_: positive Δ*T*_m_, green; negative Δ*T*_m_, orange.

### VH1-peptide interaction surfaces determined by binding affinity

We used surface plasmon resonance (SPR) to determine affinities between the pTyr EGFR peptide and the panel of VH1 mutants to further define the relationship between VH1 surface residues involved in peptide binding and subsequent dephosphorylation. The peptides were flowed over microarrays of gold surfaces with attached VH1-MBP fusion proteins for analysis of kinetic interactions by SPR. The control phosphatase, YopH, which recognizes phosphotyrosine residues regardless of sequence context (29), presented the lowest *K*_D_ for pTyr-EGFR peptide binding, and these results were consistent with a tighter binding between the two molecules in comparison to VH1. Compared to the wild-type VH1 and C110S mutant, the apparent pTyr-peptide affinities were reduced the greatest for the following VH1 mutants: K20A > K22A > M60A > T25A, T59A, as shown in Figure 4. For all of the VH1 mutants that exhibited the greatest change in *K*_D_, we noted that there were minimal protein destabilizing effects resulting from the introduction of Ala residues at these surface positions (Figure 4). Based on the SPR results, it was possible that positively charged lysine residues (Lys-20 and Lys-22) and the hydrophobic Met-60 may play a dominant role in peptide interactions.

**Figure 4.**
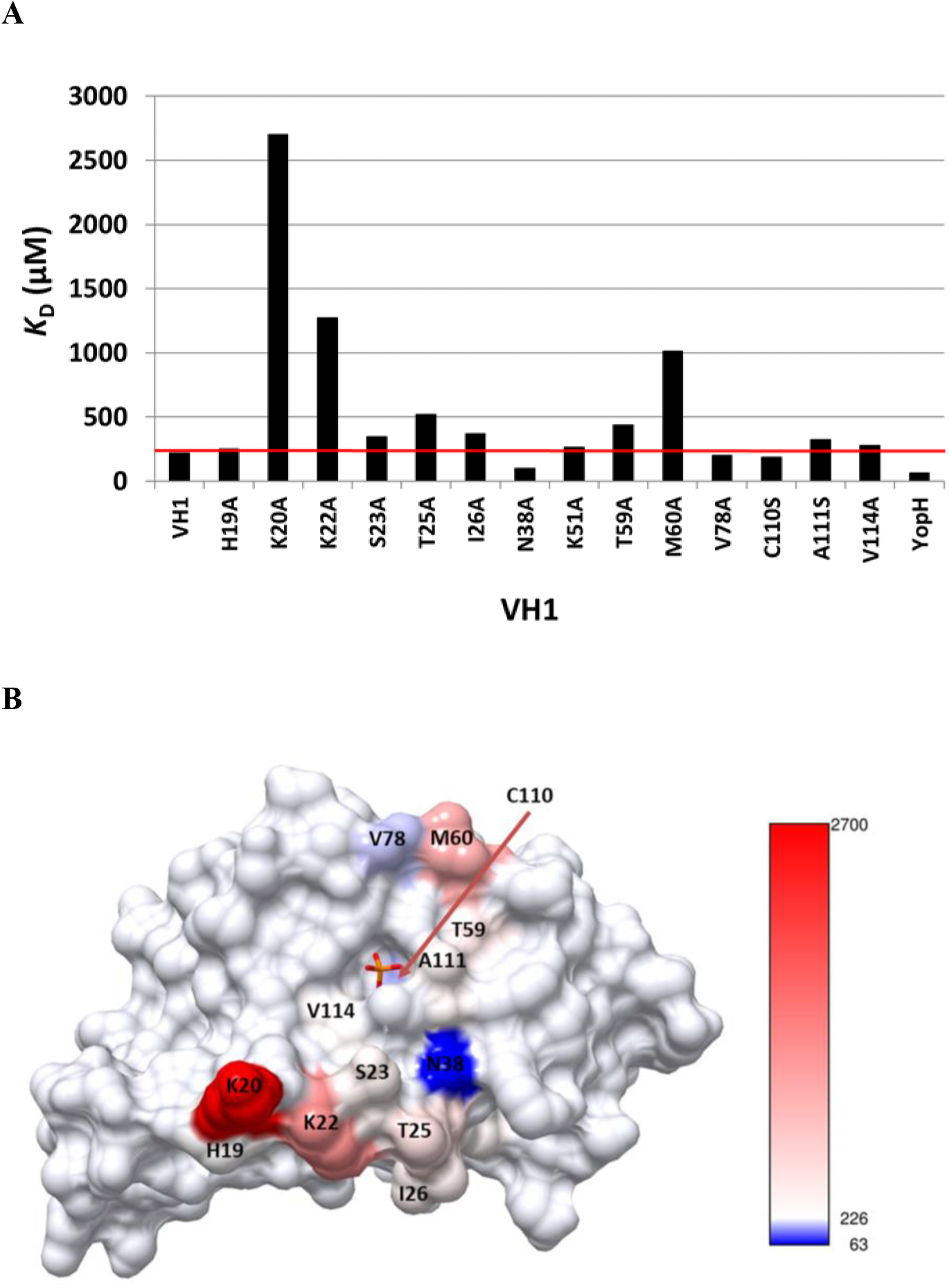
VH1 surface residues involved with phosphorylated peptide binding. **A.** EGFR peptide binding to VH1 (*K*_D_) was determined by SPR. The dashed line denotes reference K_D_ for wild-type VH1. The bacterial YopH phosphatase was used as a positive control for the experiment. **B.** Locations of VH1 surface residues by peptide affinity. The *K*_D_ for wild-type VH1 was determined to be 226 μM. Residues that have *K*_D_ values similar to the wild-type VH1 or that were not tested are colored in light-gray; residues that have a *K*_D_ ≥ wild-type VH1 are colored in red; and residues with *K*_D_ ≤ wild-type VH1 are colored in blue. The figure was generated by Chimera software (44).

To further test the hypothesis that the positively-charged residues are involved in binding to the negatively-charged EGFR peptide, we replaced lysine (K) residues on the surface of VH1 with negatively-charged aspartic acid (D). Measuring the relative distance between the phosphate molecule inside the catalytic pocket of the VH1 crystal structure and the α-carbon on the lysine residue (Figure 5A), we generated K to E mutants of four residues within 15 Å of the catalytic pocket (K20E, K22E, K62E, K87E) to accommodate hypothetical interactions with the pTyr-peptide. To evaluate catalytic interactions between the VH1 lysine mutants and peptide substrates, we constructed microarrays of biotin-labeled pTyr EGFR peptides with varying chain lengths that were spotted on streptavidin-coated nitrocellulose surfaces (Figure 5). As previously observed, dephosphorylation was greater, in general, for the longer chain length peptides. The VH1 mutants K20E, K22E, and K40E showed a small but significant decrease in peptide dephosphorylation, while K60E and K87E exhibited a minimal or negligible change in dephosphorylation compared to the wild-type VH1 (Figure 5A). Consistent with the previous kinetic peptide binding results (Figure 4), these enzymatic dephosphorylation data also indicated that residues Lys-20 and Lys-22 were involved in docking of the pTyr peptide to the VH1. To further confirm these observations, we constructed and tested the following VH1 double-mutant proteins in the immobilized dephosphorylation assay: K20E/ K22E, K20E/K62E, K20E/K87E, K22E/K62E, K22E/K87E, and K62E/K87E. Consistent with the previous results, the VH1 double mutant protein, K20E/K22E, displayed the greatest decrease in dephosphorylation of the pTyr peptides, whereas the double VH1 mutant K62E/K87E revealed no change in dephosphorylation compared to the wild-type VH1. These data strongly suggested that residues Lys-62 and Lys-87 were not involved in pTyr peptide binding. Further, all single or double lysine to glutamic acid mutants of VH1 exhibited comparable dephosphorylation of the DiFMUP molecules, suggesting that these residues have minimal effects on catalytic activity. However, the triple VH1 mutants K20E/K22E/K62E and K20E/K22E/K87E showed >50% reduction of DiFMUP dephosphorylation, indicating that introduction of three site-specific mutations may perturb the overall structure of VH1. Interestingly, we observed a greater effect on pTyr dephosphorylation with the EGFR 5-mer peptide substrate, suggesting that the aspartic acid (D) at the −4 positions of the peptides may also stabilize interactions between peptide substrate and VH1.

**Figure 5.**
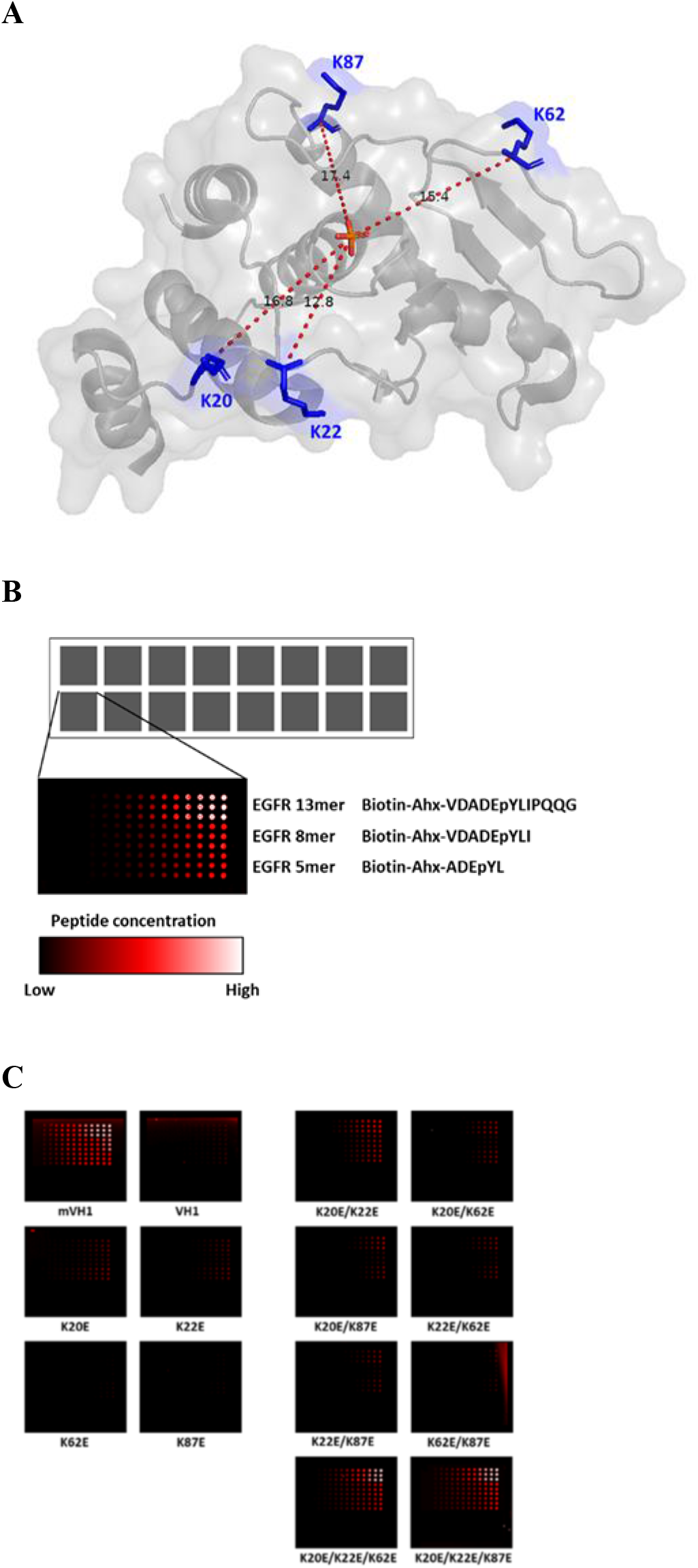
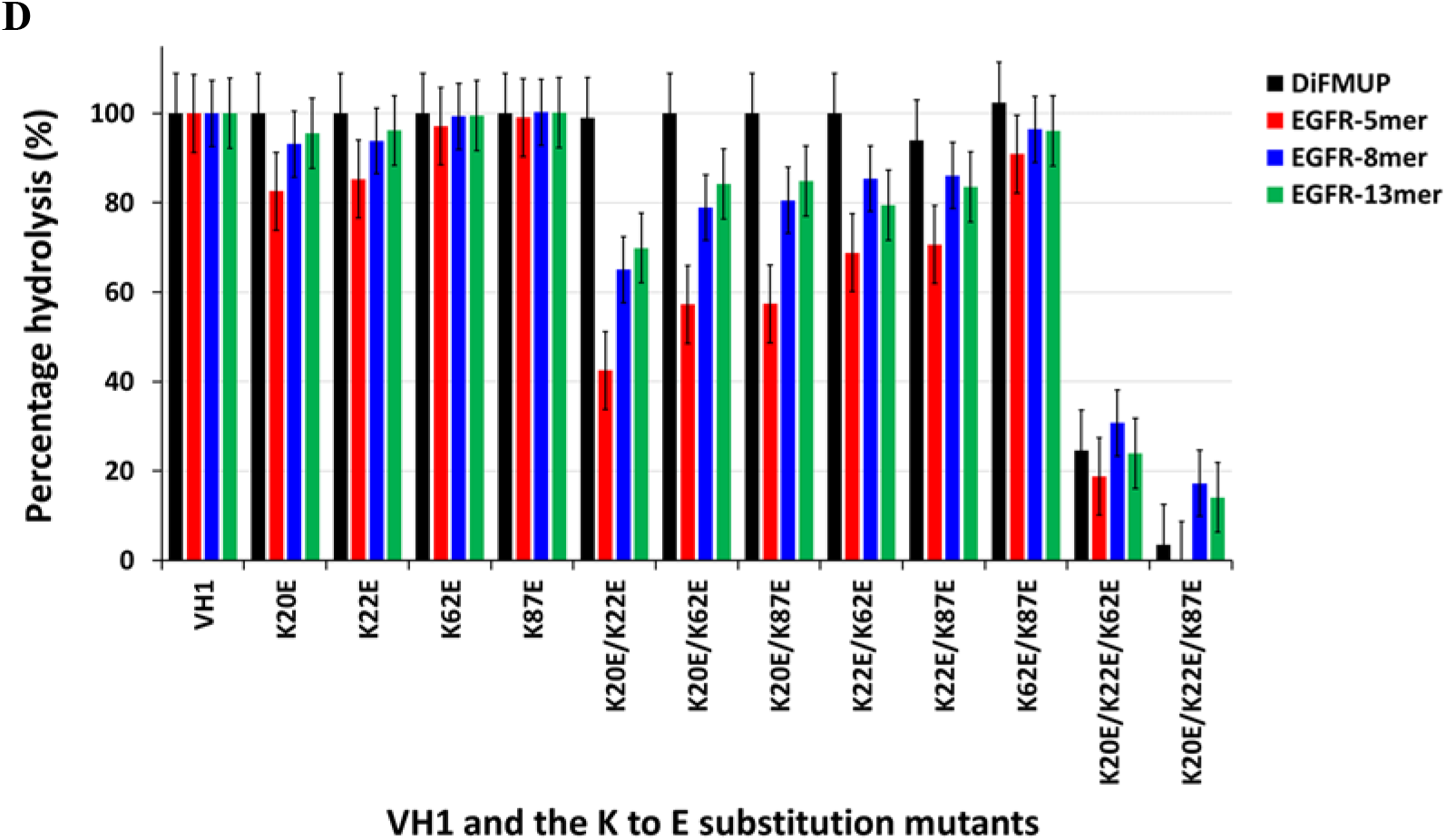
Role of VH1 surface lysine residues in peptide dephosphorylation. **A.** Locations of lysine residues on the VH1 protein surface. The distance between each of the lysine residue α-carbon atom and the phosphate atom in the catalytic site of vaccinia H1 phosphatase is ~15 Å. **B.** Representation of the peptide dephosphorylation assay. Three different lengths (11-mers, 8-mers, and 5-mers) of p-Tyr EGFR peptides were immobilized on nitrocellulose coated microscope slides in 4-400 μM spots. **C.** Scanned images of peptide dephosphorylation by K to E mutants of VH1. **D.** Summary of peptide dephosphorylation by K to E mutants of VH1. Percent dephosphorylation was calculated based on the loss of fluorescence signals compared to control.

**Figure 6.**
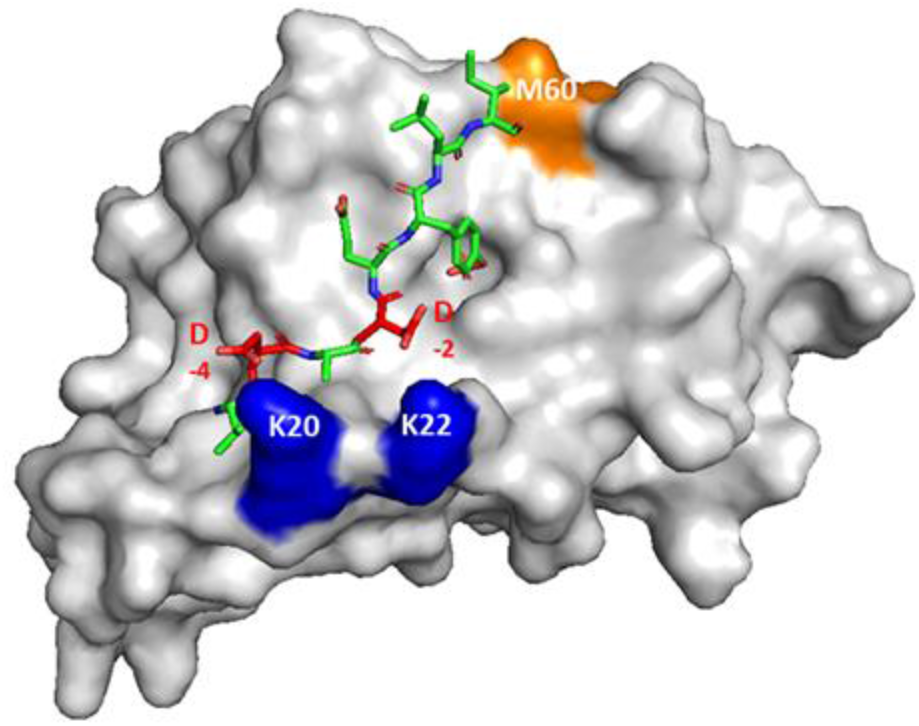
Molecular model of the pTyr EGFR peptide and VH1 phosphatase complex. The pTyr EGFR peptide is displayed as green sticks with Asp residues labeled in red. The pTyr side chain of the peptide is inserted into the catalytic pocket. VH1 Lys residues are blue and the Met residue is orange. The molecular model was generated using PyMOL software.

## DISCUSSION

We present an analysis of surface contacts between VH1 and pTyr peptide substrates that extend beyond the shallow catalytic pocket of the phosphatase. Because the VH1 active site is not readily amenable to structure-based drug development efforts, the results described here may provide a new pathway for exploring poxvirus inhibitors that target protein-protein interactions. A molecular model of pTyr peptide binding to VH1 is presented in Figure 5 to encapsulate the experimental results. In this hypothetical model, the largely positive electrostatic potential around the catalytic site orients the amino-terminus of the pTyr peptide by attracting negatively charged side chains of Asp residues at the −2 and −4 peptide positions, while the more hydrophobic carboxyl-terminus of the peptide (+1 Leu and +2 Ile positions) associates with the hydrophobic Met-60 side chain of VH1. Met-60 may also strengthen interactions by strong Van der Waals forces that are contributed by the sulfur atom (30). Presumably, the remaining key VH1 residues flanking the catalytic site that were identified in our study will further guide peptide recognition through hydrogen bonding and other mechanisms.

Our screening of potential peptide substrates identified a pTyr-EGFR peptide as an excellent substrate for studying catalytically productive interactions with VH1. It is known that engagement of EGFR increases STAT1 expression (31,32) and that poxviruses hijack EGFR-induced cell motility to spread between cells (33). Thus, it is conceivable that VH1 may dephosphorylate the endogenous EGFR during the infection cycle, though a direct linkage between the EGFR peptide sequence and poxvirus replication cycles was not established. Our molecular model considers only a single orientation for the EGFR peptide docking with VH1 and alternative configurations for other peptide sequences are possible. A study (34) of the vaccinia VH1-related (VHR) phosphatase, a mammalian cell DUSP, found that peptide substrates may bind in opposite orientations depending on the sequence motif. The amino-terminal DUSP peptide was previously reported to influence catalytic activity. We noted that mutations in the previously described dimer interface also reduced peptide dephosphorylation. In the earlier report (28), deletion of the dimerization domain helix (amino-terminal aa 1-20) greatly reduced phosphatase activity with small molecule substrates. The amino-terminal helix has also been shown to influence catalytic activity of other DUSPs and it is possible that a VH1 dimer may provide additional constraints on substrate recognition. In contrast to VH1, there is no indication that VHR dimerizes. However, it was reported that a substrate recognition segment falling outside of the VHR catalytic pocket is located between the α1-β1 residues Gly19–Pro29 (35). The α1-β1 loop of VHR surrounds the Arg-158 residue to form a positively-charged pocket for peptide binding. In a similar manner, both Lys-20 and Lys-22, which we identified as peptide-interacting residues, are enclosed by the α1-β1 loop within the vaccinia VH1 crystal structure (16). Interestingly, an analogous region that is involved in peptide substrate binding (25) was also reported for protein tyrosine phosphatase 1B (PTP1B), a negative regulator of the leptin and insulin signaling pathways. Further, and in contrast to VH1, deletion of residues 1-38 of DUSP26 has minor effect on catalytic activity (36) and the truncated protein is monomeric in solution. For DUSP26, the amino-terminal peptide chain appears to function as a scaffold for maintaining the PTP-loop conformation that is required for phosphatase activity (36).

Beyond the essential role in poxvirus infection, DUSPs are important regulators of many cellular signaling pathways. Many DUSP genes are implicated in human diseases affecting immunity and infection (37), cancer, muscle disorders, neurological (38), cardiovascular, and inflammatory diseases (39). The suit of methods described in our study may be useful for examining protein-protein interactions of these other enzyme-substrate combinations. Catalytic and kinetic interaction studies can both be accomplished by using printed microarrays of peptide substrates and mutagenized enzyme libraries as high-throughput aids for data generation. Incorporating the library of site-specific VH1 mutants into microarrays that were printed on gold-layered glass allowed us to perform multiplexed binding studies in a reusable SPR format. Previous results also demonstrated that is possible in some cases to use SPR to determine binding-on and off-rates with catalytically active phosphatases (29). Further, phosphorylated peptides reduce the complexity of protein-protein interactions to fewer testable variables compared to large protein substrates. Additional advantages are that peptides can be chemically altered with ease, kinetic as well as enzymatic activity studies can be examined, and peptides can provide structural insight for development of synthetic chemical inhibitors (40). Modification of good peptide substrates to early stage inhibitors by introduction of pTyr analogs and chemical side chains can provide progressive steps towards rational drug design (29,40,41).

## EXPERIMENTAL PROCEDURES

### Material and reagents

Dithiobis(C_2_-NTA) was purchased from Dojindo Laboratories (Kumamoto, Japan). 1-Ethyl-3-[3-dimethylaminopropyl]carbodiimide hydrochloride (EDC), *N*-hydroxysuccinimide (NHS), glycine, flexchip-blocking buffer, and ethanolamine-HCl were purchased from GE Healthcare (Piscataway, NJ, USA). NeutrAvidin was purchased from Life Technologies Corporation (Grand Island, NY, USA). The phosphorylated peptides were biotin-labeled at the N-terminus with an aminohexanoic (Ahx) acid linker and c-terminal amide group. All peptides were purchased from peptide 2.0 (Chantilly, VA, USA) or Biomatik Corporation (Wilmington, DE, USA) with at least 95% purity (sequence information in Table 1). Monoclonal anti-phospho-Tyr mouse antibody was purchased from Cell Signaling Technology, Inc. (cat. No. 9411, Beverly, MA, USA). Alexa Fluor 647 conjugated goat anti-mouse antibody was purchased from Life Technologies, Inc. (Grand Island, NY, USA). All other reagents were purchased from Sigma-Aldrich (St. Louis, MO, USA).

### Cloning, protein expression, and purification

Plasmid DNA of VH1 phosphatase from a vaccinia virus Copenhagen strain (NC_001559.1) ORFs library (42) was used as the template for PCR amplification of the full-length sequence that was cloned into pENTR/TEV/D-TOPO vector (Life Technologies Corporation, Grand Island, NY, USA). The VH1 gene was transferred into pDEST17 (Life Technologies Corporation) or pDEST-HisMBP (Addgene, Watertown, MA, USA) using LR recombination enzymes (Life Technologies Corporation, Grand Island, NY, USA) to generate GST-tagged or His-MPB-tagged VH1 respectively. Site-directed mutagenesis was used to generate various specific mutations following the QuickChange protocol (Agilent Technologies, Santa Clara, CA, USA), using overlapping primers incorporating the mutation of interest. The nucleotide sequences of all expression vectors were confirmed by sequencing. For bacterial expression, plasmids were transformed into BL21 (DE3)-plys competent cells (Life Technologies Corporation, Grand Island, NY, USA) and protein expression was induced at OD_600_ of 0.5 by adding 0.5mM IPTG. After 4 hours of induction in 30 °C, bacteria were harvested, suspended in ice-cold phosphate buffer (50 mM phosphate, 100 mM NaCl, and pH 7.4) and lysed by sonication. The lysate was centrifuged for 30 minutes at 20,000 x g and supernatant loaded on a metal chelating affinity column. The GST-VH1 recombinant protein was purified by GST affinity chromatography and His-MBP-VH1 was purified by Ni-NTA affinity chromatography (GE Healthcare, Piscataway, NJ, USA). Purified proteins were dialyzed against Bis-Tris Buffer (50mM Bis-Tris, 50 mM NaCl, 1mM EDTA, and 1mM DTT, pH 6.8).

### Multiple sequence alignment

The VH1 amino acid sequences of vaccinia virus Copenhagen (GenBank™ accession number P20495), cowpox virus Germany 91-3 (GenBank™ accession number ABD97450), monkeypox Zaire 1979-005 (GenBank™ accession number ADK39116), camelpox virus CMS (GenBank™ accession number AAG37569), and variola major virus Bangladesh-1975 (GenBank™ accession number AAA60832) were aligned with Clustal X and BLOSSOM 62 matrix. The alignment file was submitted to the ESPript server (http://espript.ibcp.fr/ESPript/ESPript/) to generate the residue alignment images.

### pTyr peptide library screening

The phosphatase substrate library contained a selection of 360 phosphorylated peptides (purchased from JPT, Berlin, Germany) that were distributed in a 384-well microtiter plate, with 250 pmol phosphorylated peptides per well. Purified VH1 phosphatase in citrate buffer (pH 6.4) was added to each well, incubated for 30 min (37 °C), and the reaction was terminated by adding malachite green assay solutions. After 15 minutes of color development, the absorbance was measured (640 nm) using a Victor plate reader (PerkinElmer, Akron, OH, USA). The free phosphate released into each well was calculated by comparison to a dilution series of phosphate standards.

### DiFMUP phosphatase assay

The DiFMUP phosphatase assay was performed using the EnzCheck phosphatase assay kit (Invitrogen) and 384-well black transparent bottom plates (Greiner). Briefly, the affinity-purified His-MBP-VH1 proteins were added to each well in Bis-Tris buffer (50mM Bis-Tris, 50mM NaCl, 0.1% Triton 100, 0.1% [w/v] BSA, 1 mM EDTA, 1 mM DTT, pH 6.5). The reaction was initiated by the addition of DiFMUP to each well and incubated for 10 min (22°C). The yield of fluorescent products from DiFMUP hydrolysis was measured with a Tecan Safire II Microplate Reader (Tecan Group Ltd., Männedorf, Switzerland), using 360nm excitation and 450nm emission wavelengths.

### Malachite green phosphatase assay

Phosphatase activities of the wild-type VH1 and mutants proteins were determined by using a malachite green assay (Update Inc., Waltham, MA, USA), following the manufacturer’s instruction. The assay was carried out in citrate buffer (0.1M Citric Acid, pH 6.4). Purified VH1 was transferred to citrate buffer by using Zeba spin desalting columns from Thermo Fisher Scientific (Waltham, MA). Phosphorylated peptides were diluted in citrate buffer and added to a 384 well plate that contained 0.5 μg/well of VH1 and 200 μM peptide in a final volume of 50 μl. After 25 minutes of incubation, the color developed by malachite green was measured at 630 nm using the Victor Spectrophotometer (Perkin Elmer). The amount of (pmol) of free phosphate released in each reaction was determined by linear regression analysis of a standard phosphate curve.

### Surface plasmon resonance microarrays

Gold coated-SPR slides and index matching oil were purchased from Horiba Scientific (Edison, NJ, USA). The SPR-PlexII (Horiba Scientific, Edison, New Jersey) instrument was cleaned and calibrated as per manufacturer’s instructions. Self-assembled monolayers (SAMs) containing the metal-chelator nitrilotriacetic acid (NTA) were formed on a gold-SPRi slide by submerging the slide in a 1mM ethanolic solution of dithiobis(C_2_-NTA) for 22 hours at 22°C. The slide was then rinsed copiously with fresh ethanol and submerged in an aqueous solution of 50 μM NiCl2 for 10 minutes. The slide was rinsed with distilled water and dried under a gentle stream of nitrogen. The His-tagged proteins (approximately 0.4 mg/mL in 50 mM Bis-Tris HCl pH 6.8) were arrayed onto the Ni(II)-coated slide in triplicate using a Continuous Flow Microspotter (CFM; Wasatch Microfluidics, Salt Lake City, Utah). After printing was complete, the proteins were covalently attached to the slide using amine-coupling chemistry. The slide was covered with a 1:1 mixture of 0.4 M EDC and 0.1 M NHS for 10 min at 22°C and then subsequently treated with 1 M ethanolamine-HCl (pH 8.5) for 15 minutes to remove any remaining NHS-esters. The slide was rinsed with distilled water and dried under a gentle stream of nitrogen.

To confirm attachment of the spotted proteins, the response signal for anti-VH1 was compared before and after a 3-minute injection of 250 mM EDTA. All kinetic experiments were performed at 25 °C with the flow set to 50 μL/min. Peptide solutions were serially diluted (500, 350, 200, 100, 50, 25, 12.5 μM) in running buffer (50 mM Bis-Tris HCl, pH 7.0) and injected in 200 μL quantities. Dissociation kinetics in running buffer were followed for about 5 minutes after the end of injections, and the surface was then regenerated with 10 mM glycine-HCl (pH 2.0). Data analysis was performed with SPRi-analysis V1.2 and ScrubberGen 2.0 software (GenOptics, Orsay, France). SPR curves were reference-subtracted from the His-MBP spots and data were processed for best fit assuming a simple 1:1 Langmuir association model for the on-rate (*k*_a_), off-rate (*k*_d_), and dissociation constant (*K*_D_) calculations. His-YopH, MBP was purchased from ProSpec Bio (ProSpec Bio, East Brunswick, New Jersey)

### Thermal shift assay

Stability of VH1 proteins was determined with a Protein Thermal Shift Dye kit (Life Technologies Corporation, Grand Island, NY, USA) following the manufacturer’s instruction. Briefly, the purified VH1 proteins were buffer exchanged with Bis-Tris buffer (pH 6.8) containing 1mM EDTA, 1mM DTT, and dispensed (0.05mg/ml and the final volume of 20μl/well) into a 96-well PCR plates. The PCR plates were sealed with Optical Quality Sealing film (BioRad, Hercules, CA, USA). The temperature was varied 25 - 95 °C in 0.5 °C increments by using an iCycle IQ5 real-time PCR system (BioRad). Relative fluorescence was measured using excitation and emission wavelengths of 580 nm and 623 nm, respectively. The protein melting temperature (*T*_m_), was obtained by determining the inflection point from the negative first derivative curve of the change in fluorescence (-d(RFU/dT), in which d(RFU) is the change in the relative fluorescence unit (RFU) and dT is the change in temperature (T). The thermal shift (Δ*T*_m_) was determined by calculating the difference between the *T*_m_ of the wild type and mutant VH1. All melting curves were the average of three experiments performed with two or more independent protein preparations.

### Phosphopeptide microarray dephosphorylation assay

The peptide dephosphorylation experiment was described previously (29), with minor changes. Briefly, biotinylated phosphorylated peptides (5 mM) were coupled with neutravidin in citrate buffer containing 40% glycerol, pH 6.4. Dilutions of peptides (500, 400, 300, 200, and 100 μM) were spotted on nitrocellulose-coated glass slides by using an ArrayJet Microarryer Marathon Argus (Arrayjet Ltd, Roslin, UK) and dried 12h under vacuum conditions. For the dephosphorylation assay, slides were first blocked with Biacore 1X Flexchip blocking buffer for 30 minutes, then washed with 1X TBS 0.2% Tween-20. While slides are still wet, a 16-well gasket cover was used to separate each well. VH1 was added (0.5μg) to each well (100μl) and incubated for 10 min (22°C). Gaskets were removed and the slides were washed with 1X TBS-0.2% Tween-20 for 30 min. To detect pTyr dephosphorylation, anti-pTyr antibody (1:1000 dilution) and goat anti-mouse IgG Alex 647 were added sequentially to slides, with wash step in between. Slides were washed with distilled water for 30 minutes, dried and scanned with a Genepix 6000B scanner (Molecular Devices, Sunnyvale, CA, USA). Data were analyzed with GenepixPro software. The percentage inhibition was calculated using the equation below:

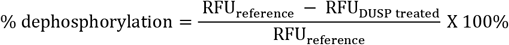

### Protein molecular modeling

The crystal structure coordinates of VH1 (PDB: 3CM3) were used to generate the surface representation of the protein structure and calculate molecular distances with PyMOL software (https://pymol.org/). To color surface residues based on experimental data, the residue attribute value was first assigned to each residue based on the experiment Kd or Δ*T*_m_. The residue attribute data were then imported into Chimera (https://www.cgl.ucsf.edu/chimera/) to generate models of surface residues with varying colors and color intensity. The molecular model of EGFR peptide binding to VH1 was generated with PyMOL software.

## ACKNOWLEDGMENTS

The contents of this publication do not necessarily reflect the views or policies of the U.S. Army, nor does the mention of trade names, commercial products or organizations imply endorsement by the U.S. Government.

## REFERENCES

1. Jezek, Z., Grab, B., Szczeniowski, M., Paluku, K. M., and Mutombo, M. (1988) Clinico-epidemiological features of monkeypox patients with an animal or human source of infection. Bull World Health Organ 66, 459–464

2. Berche, P. (2001) The threat of smallpox and bioterrorism. Trends Microbiol 9, 15–18

3. Lin, S., and Broyles, S. S. (1994) Vaccinia protein kinase 2: a second essential serine/threonine protein kinase encoded by vaccinia virus. Proc Natl Acad Sci U S A 91, 7653–7657

4. Rempel, R. E., Anderson, M. K., Evans, E., and Traktman, P. (1990) Temperature-sensitive vaccinia virus mutants identify a gene with an essential role in viral replication. J Virol 64, 574–583

5. Van Vliet, K., Mohamed, M. R., Zhang, L., Villa, N. Y., Werden, S. J., Liu, J., and McFadden, G. (2009) Poxvirus proteomics and virus-host protein interactions. Microbiol Mol Biol Rev 73, 730–749

6. Patterson, K. I., Brummer, T., O’Brien, P. M., and Daly, R. J. (2009) Dual-specificity phosphatases: critical regulators with diverse cellular targets. Biochem J 418, 475–489

7. Liu, K., Lemon, B., and Traktman, P. (1995) The dual-specificity phosphatase encoded by vaccinia virus, VH1, is essential for viral transcription in vivo and in vitro. J Virol 69, 7823–7834

8. Wishart, M. J., and Dixon, J. E. (2002) PTEN and myotubularin phosphatases: from 3-phosphoinositide dephosphorylation to disease. Trends Cell Biol 12, 579–585

9. Liu, Y., and Bankaitis, V. A. (2010) Phosphoinositide phosphatases in cell biology and disease. Prog Lipid Res 49, 201–217

10. Hunter, T. (1998) The role of tyrosine phosphorylation in cell growth and disease. Harvey Lect 94, 81–119

11. Soulsby, M., and Bennett, A. M. (2009) Physiological signaling specificity by protein tyrosine phosphatases. Physiology (Bethesda) 24, 281–289

12. Tonks, N. K., and Neel, B. G. (1996) From form to function: signaling by protein tyrosine phosphatases. Cell 87, 365–368

13. Vidovic, D., and Schurer, S. C. (2009) Knowledge-based characterization of similarity relationships in the human protein-tyrosine phosphatase family for rational inhibitor design. J Med Chem 52, 6649–6659

14. Guan, K. L., Broyles, S. S., and Dixon, J. E. (1991) A Tyr/Ser protein phosphatase encoded by vaccinia virus. Nature 350, 359–362

15. Najarro, P., Traktman, P., and Lewis, J. A. (2001) Vaccinia virus blocks gamma interferon signal transduction: viral VH1 phosphatase reverses Stat1 activation. J Virol 75, 3185–3196

16. Koksal, A. C., Nardozzi, J. D., and Cingolani, G. (2009) Dimeric quaternary structure of the prototypical dual specificity phosphatase VH1. J Biol Chem 284, 10129–10137

17. Haftchenary, S., Jouk, A. O., Aubry, I., Lewis, A. M., Landry, M., Ball, D. P., Shouksmith, A. E., Collins, C. V., Tremblay, M. L., and Gunning, P. T. (2015) Identification of Bidentate Salicylic Acid Inhibitors of PTP1B. ACS Med Chem Lett 6, 982–986

18. Fang, L., Zhang, H., Cui, W., and Ji, M. (2008) Studies of the mechanism of selectivity of protein tyrosine phosphatase 1B (PTP1B) bidentate inhibitors using molecular dynamics simulations and free energy calculations. J Chem Inf Model 48, 2030–2041

19. Liu, F., Hakami, R. M., Dyas, B., Bahta, M., Lountos, G. T., Waugh, D. S., Ulrich, R. G., and Burke, T. R., Jr. (2010) A rapid oxime linker-based library approach to identification of bivalent inhibitors of the Yersinia pestis protein-tyrosine phosphatase, YopH. Bioorg Med Chem Lett 20, 2813–2816

20. Barford, D., Flint, A. J., and Tonks, N. K. (1994) Crystal structure of human protein tyrosine phosphatase 1B. Science 263, 1397–1404

21. Raran-Kurussi, S., and Waugh, D. S. (2012) The ability to enhance the solubility of its fusion partners is an intrinsic property of maltose-binding protein but their folding is either spontaneous or chaperone-mediated. PLoS One 7, e49589

22. Tropea, J. E., Cherry, S., Nallamsetty, S., Bignon, C., and Waugh, D. S. (2007) A generic method for the production of recombinant proteins in Escherichia coli using a dual hexahistidine-maltose-binding protein affinity tag. Methods Mol Biol 363, 1–19

23. Derrien, M., Punjabi, A., Khanna, M., Grubisha, O., and Traktman, P. (1999) Tyrosine phosphorylation of A17 during vaccinia virus infection: involvement of the H1 phosphatase and the F10 kinase. J Virol 73, 7287–7296

24. Zhao, B. M., Keasey, S. L., Tropea, J. E., Lountos, G. T., Dyas, B. K., Cherry, S., Raran-Kurussi, S., Waugh, D. S., and Ulrich, R. G. (2015) Phosphotyrosine Substrate Sequence Motifs for Dual Specificity Phosphatases. PLoS One 10, e0134984

25. Jia, Z., Barford, D., Flint, A. J., and Tonks, N. K. (1995) Structural basis for phosphotyrosine peptide recognition by protein tyrosine phosphatase 1B. Science 268, 1754–1758

26. Sarmiento, M., Puius, Y. A., Vetter, S. W., Keng, Y. F., Wu, L., Zhao, Y., Lawrence, D. S., Almo, S. C., and Zhang, Z. Y. (2000) Structural basis of plasticity in protein tyrosine phosphatase 1B substrate recognition. Biochemistry 39, 8171–8179

27. Zhang, Z. Y., Maclean, D., McNamara, D. J., Sawyer, T. K., and Dixon, J. E. (1994) Protein tyrosine phosphatase substrate specificity: size and phosphotyrosine positioning requirements in peptide substrates. Biochemistry 33, 2285–2290

28. Koksal, A. C., and Cingolani, G. (2011) Dimerization of Vaccinia virus VH1 is essential for dephosphorylation of STAT1 at tyrosine 701. J Biol Chem 286, 14373–14382

29. Hogan, M., Bahta, M., Cherry, S., Lountos, G. T., Tropea, J. E., Zhao, B. M., Burke, T. R., Jr., Waugh, D. S., and Ulrich, R. G. (2013) Biomolecular Interactions of small-molecule inhibitors affecting the YopH protein tyrosine phosphatase. Chem Biol Drug Des 81, 323–333

30. Gomez-Tamayo, J. C., Cordomi, A., Olivella, M., Mayol, E., Fourmy, D., and Pardo, L. (2016) Analysis of the interactions of sulfur-containing amino acids in membrane proteins. Protein Sci 25, 1517–1524

31. Cheng, C. C., Lin, H. C., Tsai, K. J., Chiang, Y. W., Lim, K. H., Chen, C. G., Su, Y. W., Peng, C. L., Ho, A. S., Huang, L., Chang, Y. C., Lin, H. C., Chang, J., and Chang, Y. F. (2018) Epidermal growth factor induces STAT1 expression to exacerbate the IFNr-mediated PD-L1 axis in epidermal growth factor receptor-positive cancers. Mol Carcinog 57, 1588–1598

32. Han, W., Carpenter, R. L., Cao, X., and Lo, H. W. (2013) STAT1 gene expression is enhanced by nuclear EGFR and HER2 via cooperation with STAT3. Mol Carcinog 52, 959–969

33. Beerli, C., Yakimovich, A., Kilcher, S., Reynoso, G. V., Flaschner, G., Muller, D. J., Hickman, H. D., and Mercer, J. (2019) Vaccinia virus hijacks EGFR signalling to enhance virus spread through rapid and directed infected cell motility. Nat Microbiol 4, 216–225

34. Luechapanichkul, R., Chen, X., Taha, H. A., Vyas, S., Guan, X., Freitas, M. A., Hadad, C. M., and Pei, D. (2013) Specificity profiling of dual specificity phosphatase vaccinia VH1-related (VHR) reveals two distinct substrate binding modes. J Biol Chem 288, 6498–6510

35. Yuvaniyama, J., Denu, J. M., Dixon, J. E., and Saper, M. A. (1996) Crystal structure of the dual specificity protein phosphatase VHR. Science 272, 1328–1331

36. Won, E. Y., Lee, S. O., Lee, D. H., Lee, D., Bae, K. H., Lee, S. C., Kim, S. J., and Chi, S. W. (2016) Structural Insight into the Critical Role of the N-Terminal Region in the Catalytic Activity of Dual-Specificity Phosphatase 26. PLoS One 11, e0162115

37. Lang, R., and Raffi, F. A. M. (2019) Dual-Specificity Phosphatases in Immunity and Infection: An Update. Int J Mol Sci 20

38. Bhore, N., Wang, B. J., Chen, Y. W., and Liao, Y. F. (2017) Critical Roles of Dual-Specificity Phosphatases in Neuronal Proteostasis and Neurological Diseases. Int J Mol Sci 18

39. Pulido, R., and Hooft van Huijsduijnen, R. (2008) Protein tyrosine phosphatases: dual-specificity phosphatases in health and disease. FEBS J 275, 848–866

40. Hogan, M., Bahta, M., Tsuji, K., Nguyen, T. X., Cherry, S., Lountos, G. T., Tropea, J. E., Zhao, B. M., Zhao, X. Z., Waugh, D. S., Burke, T. R., Jr., and Ulrich, R. G. (2019) Targeting Protein-Protein Interactions of Tyrosine Phosphatases with Microarrayed Fragment Libraries Displayed on Phosphopeptide Substrate Scaffolds. ACS Comb Sci 21, 158–170

41. Burke, T. R., Jr., Ye, B., Yan, X., Wang, S., Jia, Z., Chen, L., Zhang, Z. Y., and Barford, D. (1996) Small molecule interactions with protein-tyrosine phosphatase PTP1B and their use in inhibitor design. Biochemistry 35, 15989–15996

42. Schmid, K., Keasey, S. L., Pittman, P., Emerson, G. L., Meegan, J., Tikhonov, A. P., Chen, G., Schweitzer, B., and Ulrich, R. G. (2008) Analysis of the human immune response to vaccinia by use of a novel protein microarray suggests that antibodies recognize less than 10% of the total viral proteome. Proteomics Clin Appl 2, 1528–1538

43. Robert, X., and Gouet, P. (2014) Deciphering key features in protein structures with the new ENDscript server. Nucleic Acids Res 42, W320–324

44. Pettersen, E. F., Goddard, T. D., Huang, C. C., Couch, G. S., Greenblatt, D. M., Meng, E. C., and Ferrin, T. E. (2004) UCSF Chimera--a visualization system for exploratory research and analysis. J Comput Chem 25, 1605–1612

