## Supplemental figures for "Peptide recognition and dephosphorylation by the vaccinia VH1 phosphatase"

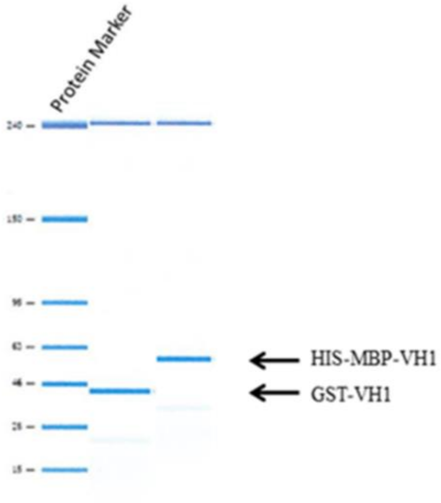


**Figure S1. SDS gel of purified GST-VH1 and HIS-MBP-VH1.** The purified GST-VH1 and His-MBP-VH1 proteins were first resolved on a 4-20% polyacrylamide gel in SDS and stained with Coomassie brilliant blue; lane A is the molecular weight standard.

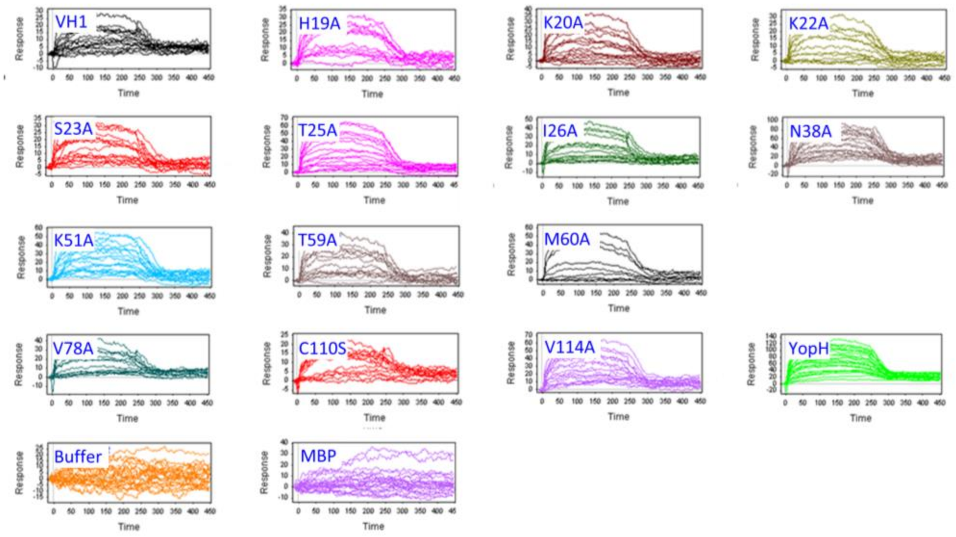


**Figure S2.** Representative sensorgrams of pTyr peptide (EGFR) interactions with VH1 site-specific mutants.

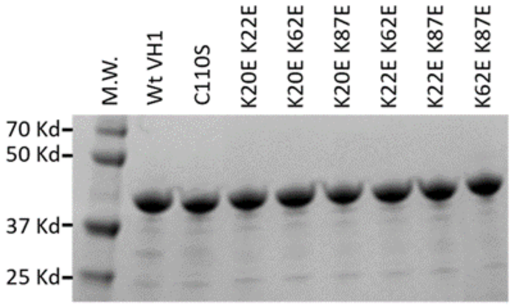


**Figure S3. Representative SDS-PAGE profiles of wild-type His-MBP-VH1 and site-directed K to E mutants.** The purified proteins were run on 10% SDS-polyacrylamide gel electrophoresis under reducing conditions. The gel was stained with Coomassie Blue.
